# Computational Re-Design of Synthetic Genetic Oscillators for Independent Amplitude and Frequency Modulation

**DOI:** 10.1101/179945

**Authors:** M. Tomazou, M. Barahona, K. Polizzi, G.-B. Stan

**Affiliations:** Department of Bioengineering, Imperial College London; Department of Mathematics, Imperial College London; Department of Life Sciences, Imperial College London; Centre for Synthetic Biology and Innovation, Imperial College London

## Abstract

Engineering robust and tuneable genetic clocks is a topic of current interest in Systems and Synthetic Biology with wide applications in biotechnology. Synthetic genetic oscillators share a common structure based on a negative feedback loop with a time delay, and generally display only limited tuneability. Recently, the dual-feedback oscillator was demonstrated to be robust and tuneable, to some extent, by the use of chemical inducers. Yet no engineered genetic oscillator currently allows for the *independent* modulation of amplitude and period. In this work, we demonstrate computationally how recent advances in tuneable synthetic degradation can be used to decouple the frequency and amplitude modulation in synthetic genetic oscillators. We show how the range of tuneability can be increased by connecting additional input dials, e.g. orthogonal transcription factors that respond to chemical, temperature or even light signals. Modelling and numerical simulations predict that our proposed re-designs enable amplitude tuning without period modulation, coupled modulation of both period and amplitude, or period adjustment with near-constant amplitude. We illustrate our work through computational re-designs of both the dual-feedback oscillator and the repressilator, and show that the repressilator is more flexible and can allow for independent amplitude and near-independent period modulation.

## 1. Introduction

Accurate temporal control of biological processes is of fundamental importance across all kingdoms of life and is often realised through genetic ‘clocks’, *i.e.,* transcriptional networks with periodic gene expression. Heart beats, cell cycles, circadian rhythms, developmental processes and energy metabolism are all, in one way or another, driven by robust genetic clocks^1-3^. Such oscillators can also form the building blocks to engineer complex genetic networks that can be used to synchronise cellular activity or to optimise the efficiency of metabolic pathways. Understanding the design principles of genetic oscillators and finding ways to engineer and precisely control them is therefore of crucial importance, as conveyed by the numerous synthetic oscillators proposed to date^4-13^.

All genetic transcriptional oscillators are based on the same principle introduced in 1963 by Goodwin^4^: a negative feedback loop with a time-delay. Perhaps the best studied example of such a system is the **repressilator** (RLT)^5^, a ring of transcriptional repressors acting in sequence, thus providing an intrinsic lag or delay. More recently, it was demonstrated that the addition of a positive feedback loop (as is commonly encountered in nature) increases the robustness and adds tuneability to the oscillator’s amplitude and period^14-16^. A prime example in this category is the **dual-feedback oscillator** (DFO)^10^, which comprises a relatively slow negative feedback loop and a faster positive feedback loop. The DFO has been shown to function robustly when implemented using different components^13^, and also as part of large networks in different organisms and conditions^17,18^ as well as in populations of cells^13,19,20^. However, an important feature yet to be implemented in either the DFO or other genetic oscillators is the ability to control the period and amplitude of oscillations *independently*. Such control would allow a wide range of applications, including frequency analysis of downstream networks^21,22^, frequency encoding and pulse-based signal processing^23^, development of two-dimensional biosensors where period and amplitude give distinct responses, designs for burden relief and metabolic pathway optimisation or even periodic administration of therapeutic molecules.

In this work, we use mathematical modelling and numerical simulation to show how simple, implementable modifications of the DFO^10^ and RTL^5^ allow the user to control their amplitude and period independently and over an increased dynamic range. Through deterministic numerical simulations of a parsimonious model, we demonstrate how to introduce orthogonal inputs or “dials”^24^ based on dual-input promoters^25,26^, that do not affect the core topology of the DFO and RLT, yet allow oscillations to be tuned in (a) amplitude only, (b) period and amplitude simultaneously, or (c) period with the amplitude maintained at near-constant levels. In addition, some of our re-designs allow the system to transition from a stable OFF state, to a tuneable oscillatory regime, and finally to a stable ON state as a function of the input dials (See SI, S6). Hence, the same architecture can be used to generate controllable oscillations and / or switching between high and low steady-states.^27^

## 2. Results

Our main objective is to re-design genetic oscillators with enhanced amplitude and frequency tuneability. We consider the two most popular oscillators in synthetic biology (DFO and RTL) and discuss key modifications of their network architecture that extend the range of achievable amplitudes and periods, at the same time decoupling these characteristics. To keep our results general, we study parsimonious models that adequately reproduce their behaviours. The models comprise three modules: (i) a core oscillator module composed of a negative feedback loop with delay mediated by repressors (with an additional positive feedback loop for the DFO); (ii) an output module, with the gene of interest as a readout; (iii) a sink module where the species of the oscillator and output modules are channelled for enzymatic degradation. Our analysis identified the post-translational coupling introduced by enzymatic degradation (a.k.a. protease ‘queuing effect’^28^) as a major challenge for independent amplitude modulation. This coupling affects the degradation rate of the transcription factors (TF) and consequently the period^28,29^. To mitigate this effect, we introduce an orthogonal enzymatic degradation pathway^30^ that enables modulation of amplitude with no impact on the period.

To achieve independent period modulation, we followed two approaches. First, a re-design that modulates the expression levels of the protease showed that the oscillators exhibit a significant increase in the achievable range of periods. Second, we considered tuning of the negative feedback loop by modulating the expression rate of one of the repressors in the RTL. Our simulations showed that the period can be tuned within a greater range compared to chemical induction alone, while the amplitude is kept nearly constant.

Below we give a description of the original and modified networks considered in this work. Our coarse-grained phenomenological models employ Hill-type functions^31^ to capture the regulatory action of single and dual-input promoters as well as the inputs of DFO and RLT. Although these models are too simplistic to reproduce the full range of dynamics captured by detailed mechanistic models using characterised biological parts^10^, they capture the core responses of the oscillators and produce stable oscillations with periods within the range observed in experimentally implemented systems^5,10^.

### 2.1. The dual-feedback oscillator and its redesigns

#### 2.1.1 The original DFO

The DFO basic structure (Fig. 1A) comprises a positive and a negative feedback loop mediated by a transcriptional activator (A) and repressor (R), respectively. The output is defined as the oscillating protein of interest denoted by G. All three proteins (A, R, and G) are tagged for fast enzymatic degradation *via* the same protease (C). Note that a basic difference in our modelled network from the original published DFO model^10^, is that we take in account the output protein and its queuing effect^28^ on the degradation rates.

**Figure 1.**
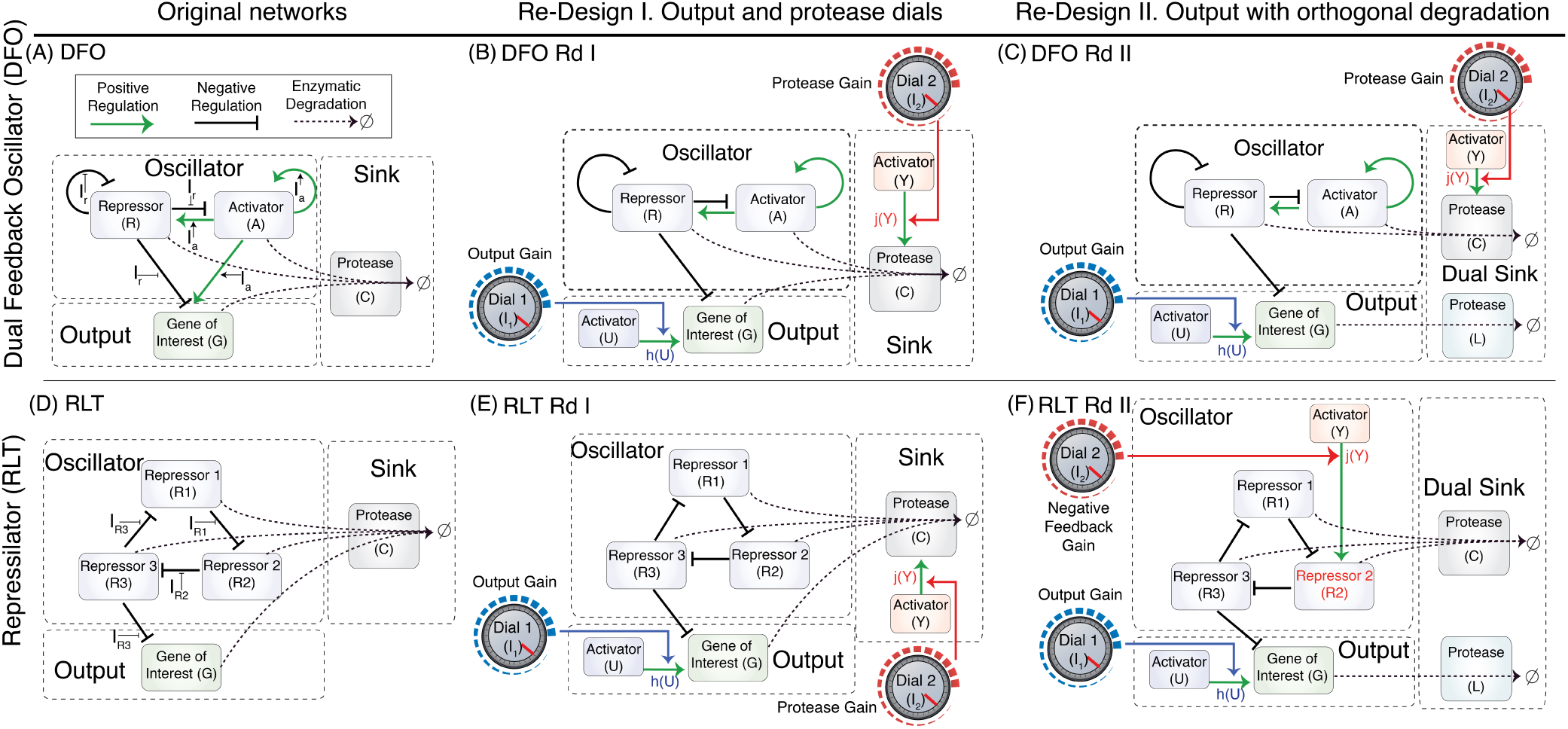
Considered architectures of the dual-feedback oscillator (**A**,**B**,**C**) and repressilator (**D,E,F**). For each architecture, positive (green arrow lines) and negative (black solid lines) regulations are indicated. Enzymatic degradation reactions are indicated by black dashed arrows. Each network consists of a core oscillator module, a degradation (sink) module, and an output module. The inputs considered for the original DFO and RLT designs are inducers titrating the effect of the TFs in the oscillator module. For all the redesigned networks (labelled Rd), an input *I*_*1*_ (Dial 1) controls the affinity of an orthogonal TF *U*, which in turn controls the expression rate of the gene of interest (*G*) in the output module. For the redesigns DFO Rd I and the RLT Rd I, a second external input *I*_*2*_ (Dial 2) is used to modulate the levels of the protease *C via* TF *Y.* For redesigns DFO Rd II and RLT Rd II, a second orthogonal protease *L* is used to decouple the degradation of *G* from the degradation of the TF in the oscillator module. In RLT Rd II, instead of modulating the amount of *C* like in DFO Rd II, we use the second input *I*_*2*_ to modulate the expression rate of the repressor *R2*.

The rate of change of the mRNA concentrations of the three genes *m*_*i*_ is given by:

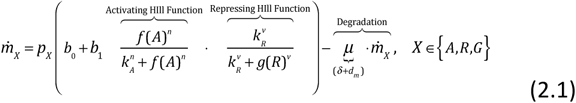

where *p*_*X*_ is the gene copy number, and *b*_*0*_ and *b*_*1*_ the basal and maximum transcription rate constant, respectively. *k*_*A*_ and *k*_*R*_ are Hill constants describing the amounts of transcription factors (TFs) required for half-maximal activation and repression rates, respectively, whereas *n* and *v* are Hill cooperativities reflecting whether the TFs are active as monomers or multimers. As per the original published model for DFO^10^ we used higher cooperativity degree for the repressor (*v*=4) compared to the activator (*n*=2). The removal rate constant per mRNA is given by *μ* and captures the sum of the dilution rate due to cell division (*δ*) and the mRNA decay rate (*d*_*m*_). The inducer-dependent functions *f(A)* and *g(R)* are defined as:

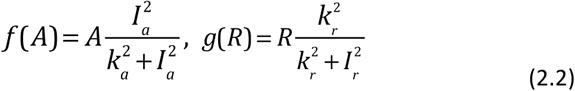

where *I*_*a*_ and *I*_*r*_ are the inducer concentrations that increase or decrease the effect of the activator (A) or repressor (R) with half-maximal constants *k*_*a*_ and *k*_*r*_, respectively. For simplicity, their Hill cooperativities are set to 2 to represent a typical dimeric transcription factor. Additional simulation results (not shown here) indicate that similar qualitative behaviours are obtained when considering other Hill cooperativities, e.g. cooperativities 1 and 4 in (2.2). In what follows, we therefore only present simulation results for Hill cooperativities set to 2.

The dynamics of the associated proteins (folded and unfolded) are described as follows:

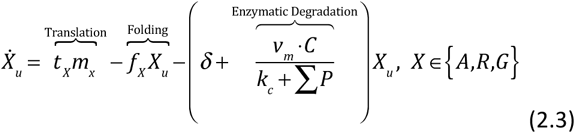

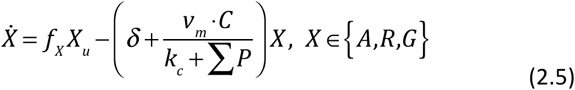

where *X*_*u*_ and *X* are the concentrations of unfolded and folded proteins, respectively. Here *t*_*X*_ and *f*_*X*_ are the translation and folding rate constants, respectively, and *δ* is the dilution rate due to cell division. The enzymatic degradation term obeys Michaelis-Menten (MM) kinetics^28^ with *v*_*m*_ representing the catalysis rate, *k*_*c*_ its MM constant and *C* the total concentration of protease in the cell. ∑*P* represents the total concentration of degradation tags competing for the same protease sites^28^, denoted hereafter as ‘the load’. Assuming TFs with identical degradation tags can bind on the protease binding site with equal affinities then

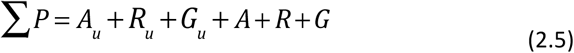

The above model was parametrised to reproduce the oscillatory behaviour observed in experiments^5,10^. The full set of equations, kinetic scheme, assumptions, and parameter values is given in SI (S7).

#### 2.1.2 Dual-feedback oscillator re-designs

DFO features an output (*G*) whose genetic expression is controlled by a repressor (*R*), so that *G* is periodically expressed when *R* levels periodically transition from high to low during an oscillation. Through numerical simulations and analysis of the Ordinary Differential Equation (ODE) model (2.1)-(2.5), we explore computationally a number of re-designs of the basic DFO architecture. These redesigns involve the introduction of additional biological dials inserted at different locations in the DFO gene regulation network (Fig. 1B, C). The external inputs for tuning the oscillation are chemical inducers (*I*_*a*_ and *I*_*r*_) that modulate the affinity of the TFs to their cognate promoter binding sites, and, therefore, their transcriptional regulation effectiveness.

##### Re-design 1: DFO Rd I

The first re-design DFO Rd I (Fig. 1B) introduces an activation input (*U*) that controls the expression of G orthogonally to the TF (R) of the core oscillator module. This can be achieved by a dual-input promoter^25,26^ that responds to two different TFs (*U* and *R*). The total amount of *U* is assumed to be unregulated (constitutive gene expression) and constant at steady state, but the activation effect of *U* on *G* is modulated by an external input signal *I*_*1*_. In this re-design, the protease expression rate is also tuneable *via* an orthogonal TF (*Y*). The effect of *Y* on *C* is modulated by an external input signal *I*_*2*_.

The corresponding model is thus updated as follows:

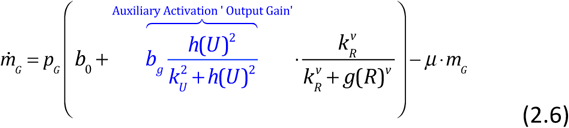

where the affinity of the activating TF *U* is modulated by the input (inducer) *I*_*1*_ which is orthogonal to *I*_*a*_ and *I*_*r*_. The function *h(U)* in (2.6) captures the influence of *I*_*1*_ on the steady state concentration of *U*, captured with a Hill function with cooperativity *2* and half-activation constant *k*_*1*_:

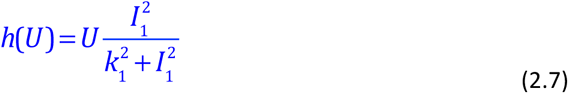

The dynamics of *G* allows the output to oscillate following oscillations in *R*, and the rate of expression of *G* when *R* is not fully repressing is titrated by *I*_*1*_. As in (2.2) using cooperativity exponents other than 2 gives similar results. In addition, the steady state levels of *C* vary as a function of a second input *I*_*2*_, as follows:

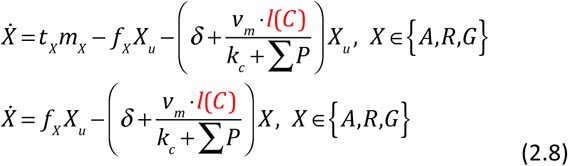

where the protease concentration *C* is a function of *Y* modulated by the second input, *I*_*2*_:

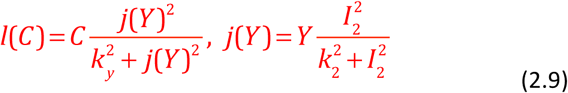

##### Re-design 2: DFO Rd II

The second re-design DFO Rd II (Fig. 1C) differs from Rd I in the degradation mechanism. In DFO Rd II, an additional orthogonal protease (*L*) is dedicated to the specific degradation of the output (*G*). This second enzymatic degradation pathway eliminates the post-translational coupling between *G* and the TFs of the core oscillator module, *i.e.,* TFs *A,R* are targeted for degradation by *C*, whereas *G* is specifically targeted for degradation by the orthogonal protease *L*^30^:

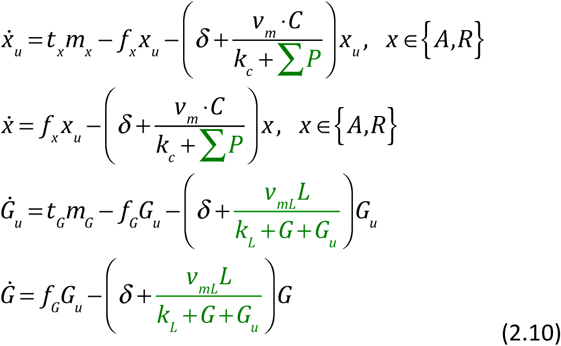

where *v*_*mL*_ and *k*_*L*_ are the Michaelis-Menten parameters for the secondary enzymatic degradation pathway. The load for the primary degradation pathway is now independent of the output (*G*)

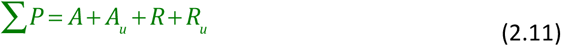

### 2.2 Computational characterisation of the DFO re-designs

The experimental implementation of the original DFO (Fig. 1A) employed two of the most commonly used TFs, LacI and AraC, which can be controlled by the chemical inducers IPTG and arabinose, respectively. We model the effect of these inducers by varying the parameters *I*_*a*_ *and I*_*r*_. Fig. 2A, B shows model predictions of how the amplitude and period change as a result of varying the concentrations of the inputs *I*_*a*_, *I*_*r*_, *I*_*1*_ and *I*_*2*_ from 0 to saturating levels with respect to their corresponding Hill functions (2.2), (2.7) and (2.9).

**Figure 2.**
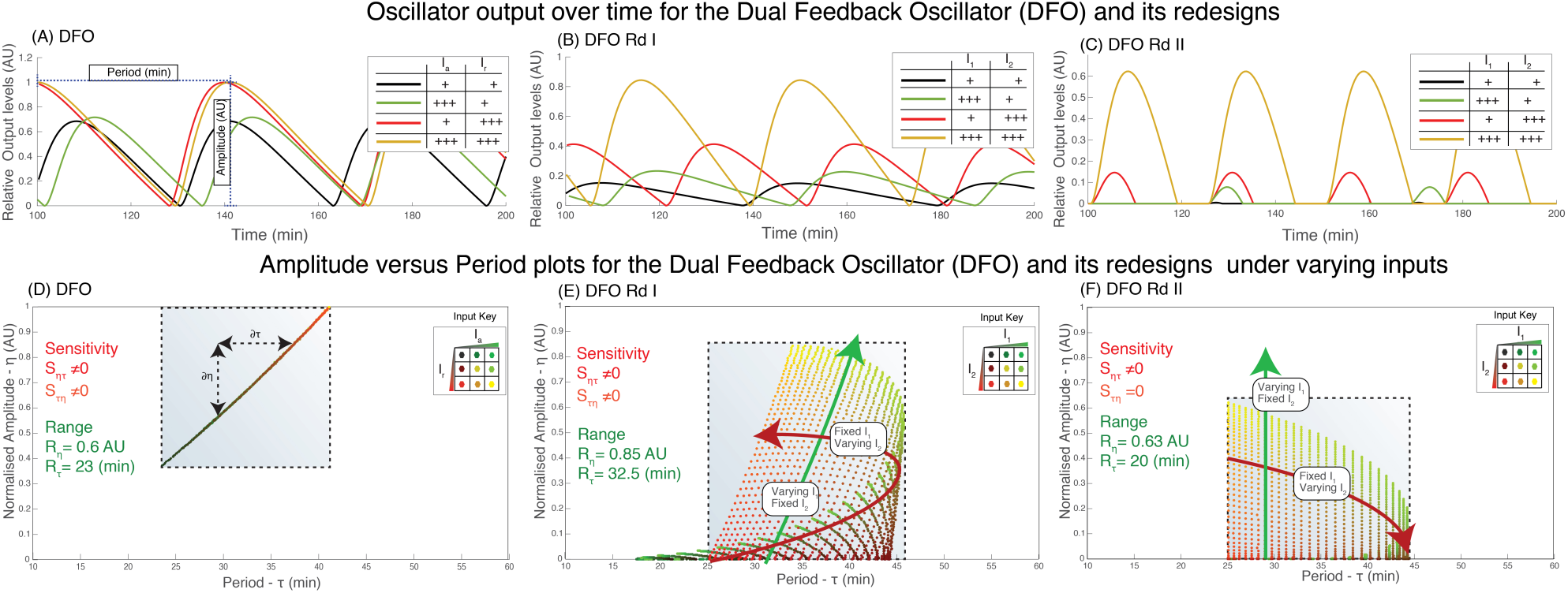
**(A,B,C)** Time course simulations showing the oscillatory output *G* over time under three different induction conditions (+ = 20 uM, +++ = 100 uM). In DFO (A) and DFO Rd I (B) when an input results in an increase in the amplitude, the period is elongated. In DFO Rd II the amplitude can be increased with no effect on the period (τ=40 min for black and green and τ=25 min for red and orange trajectories). *I*_*2*_ primarily affects the period but with some impact on the amplitude. **(D,E,F)** Amplitude versus period dot plots for DFO, DFO Rd I and DFO Rd II, respectively. The colour of the dot indicates the amount of input with the green component indicating *I*_*1*_ and the red component indicating *I*_*2*_. Yellow indicates that both inputs are high while black corresponds to their absence. (D) The original DFO exhibited a narrower range for both amplitude and period and either affects both characteristics in a highly-coupled manner. (E) The DFO Rd I shows a wider tuneable range with *I*_*1*_ primarily increasing the amplitude and *I*_*2*_ modulating the period. However, none of these characteristics were independently tuneable. (F) Finally, in the DFO Rd II the amplitude can be tuned in response to *I*_*1*_ independently of the period. The opposite was not feasible in this case.

The simulation results show that period and amplitude are correlated with *I*_*a*_ and *I*_*r*,_ in the original DFO (Fig 2A,D). Hence both inputs simultaneously affect amplitude and period so that these characteristics cannot be independently tuned. This is quantified through the local cross-sensitivities for a given amplitude and period value *(η*^*\**^, *τ*^*\**^) defined as:

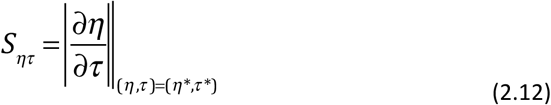

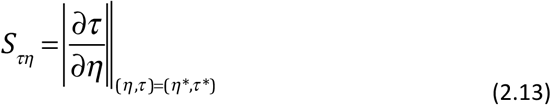

The cross-sensitivities for the entire range of amplitudes and periods are presented in SI (Fig. S1).

#### 2.2.1 DFO Rd I exhibits an increased range of achievable amplitudes and periods

Our simulations show that the range of period and amplitude tuneability of DFO Rd I is increased (Fig. 2B, E). However, although the amplitude can be tuned over a wider range, the increase in amplitude is accompanied by a nonlinear shift in the period. This is explained by the post-translational coupling between the output *G* and the TFs in the core oscillatory module introduced by the protease queuing effect, which results from the sharing of the protease among the total amount of degradation tags in the system. Hence, the degradation for G and all TFs becomes slower, which in turn increases the period. To investigate whether this effect can be attenuated by increasing the abundance of protease we use a second external input *I*_*2*_ to modulate the expression rate of the protease *C* (*via* the additional TF, *Y*). Fig. 2E shows that even with the highest levels of *C* (red points) the period is still affected by the increase in amplitude when *I*_*1*_ is increased from 0 (red dots) to saturating amounts (yellow dots).

Interestingly, modulating the amount of the protease increases the range of achievable periods by about 70% compared to the original DFO. At moderate to high amounts of protease, the sensitivity of the amplitude is relatively small but increases as the protease drops to low levels.

#### 2.2.2 DFO Rd II exhibits independent tuneability of period and amplitude

Our objective is to find oscillatory re-designs and associated conditions for which the cross-sensitivities (2.12-2.13) are significantly reduced. In DFO Rd II we aimed to eliminate the observed post-translational coupling by introducing an additional, orthogonal protease dedicated to the degradation of the output G. The results in Fig. 2C, F show a negligible sensitivity of the period to changes in the amplitude (S_τη_=0) over the full range of accessible amplitudes, *i.e.,* the amplitude is tuneable over the full range with no effect on the period. However, the amplitude to period sensitivity S_ητ_ was increased compared to DFO Rd I. A downside of DFO Rd II is a range of achievable periods comparable to that of the original DFO, although, for a given period, an extended range of amplitudes is accessible by varying *I*_*1*_.

### 2.3. The Repressilator re-designs

#### 2.3.1. The original RLT

The original repressilator model comprises a ring of three TFs, each repressing the following gene in the ring (Fig. 1D). Compared to the original RLT^5^, we model the enzymatic degradation mechanism catalysed by the protease *C*, and we include explicitly the output gene module (*G*). In addition, the repression is modulated by external inputs *I*_*Ri*_, with *i* ∊{1,2,3}.

Similarly to above, we derived a model for RLT, as follows. The mRNA rate equation is modelled as:

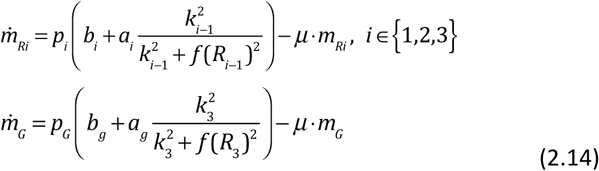

where,

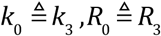

and the induction function for each inducer is given as:

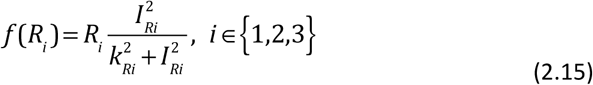

The unfolded and folded protein dynamics are given by:

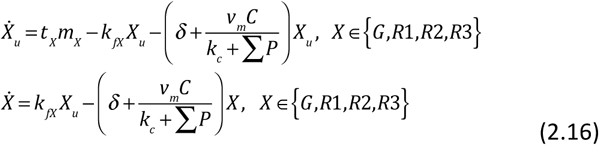

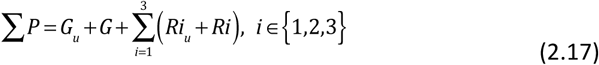

#### 2.3.2. Repressilator re-designs

The redesigns of RLT follow similar lines to the ones presented for DFO, yet the additional flexibility supported by the RLT architecture leads to more independent amplitude and period tuning.

##### Re-design 1: RLT Rd I

Two orthogonal “dials” (TFs *U* and *Y*) are introduced in RLT Rd I (Fig. 1E): *U* modulates the expression of *G* a*s* a function of the external input *I*_*1*,_ whereas *Y* modulates the expression of the degradation protease *C* as a function of the external input *I*_*2*_.

Following the same structural modifications discussed for DFO Rd I, the output mRNA rate equation is given by:

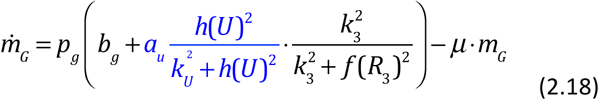

with *h(U)* given in (2.7), and the second input *I*_*2*_ regulating the amount of protease in the system:

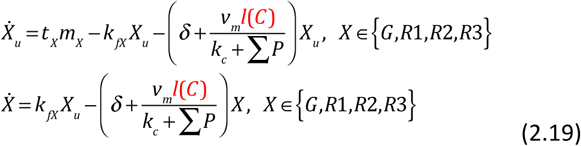

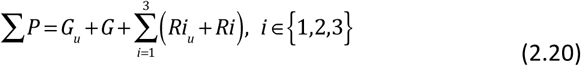

where *l(C)* is

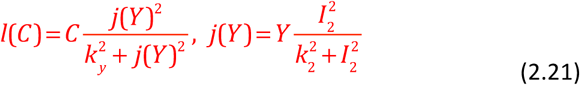

##### Re-design 2: RLT Rd II

In the second redesign (Fig. 1F), *R1* and *R3* obey the same regulation scheme as in RLT Rd I, while *R2* is now regulated by a dual-input promoter that is repressed by *R1* and activated by *Y*:

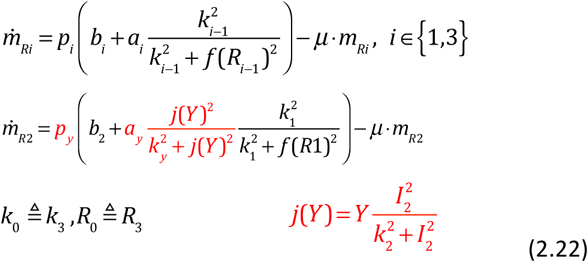

Furthermore, a secondary protease *L* is specifically dedicated to the enzymatic degradation of the output gene of interest (*G*) yielding:

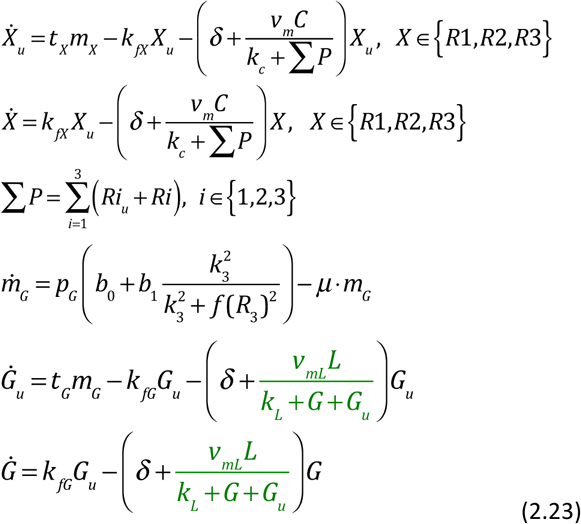

### 2.4 Computational characterisation of the RLT re-designs

The numerical simulations of the original RLT (Fig. 3A, D) demonstrate the effect of the inputs *I*_*R2*_ and *I*_*R3*_ corresponding to chemical inducers used to modulate the affinity of repressors. Other combinations of titratable inputs (*I*_*R1*_*-I*_*R2*_ *and I*_*R1*_*-I*_*R3*_) shown in SI (Fig. S2) produced similar results. The period is predicted to be tuneable over a range of approximately 185 min, whereas the amplitude range is narrower. As for the original DFO, the amplitude and period are highly correlated; hence tuning the amplitude and the period of the RLT independently is not possible.

**Figure 3.**
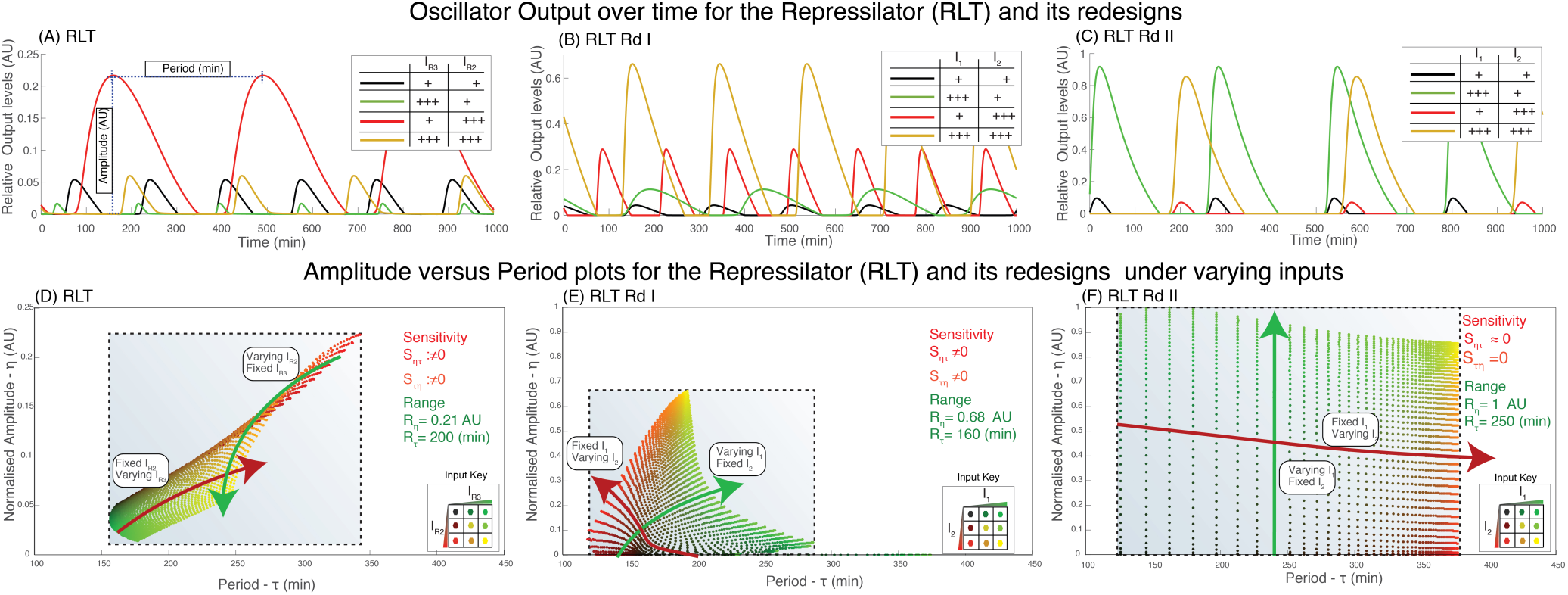
**(A,B,C)** Time course simulations of the output *G* over time under four different input conditions (+ = 20 uM, +++ = 100 uM). Similar to DFO and DFO Rd I, numerical simulation results show that the period of RLT in **(D)** and RLT Rd I in **(E)** is significantly sensitive to input *I*_*1*_ such that the amplitude cannot be modulated independently of the period. On the other hand, numerical simulations presented in **(F)** indicate that the re-design corresponding to RLT Rd II allows to tune the amplitude of the oscillations (green trajectory) over a wider range than both RLT and RLT Rd I and with no effect on the period. More surprising, tuning the period by modulating *I*_*2*_, had a minimal effect on the amplitude. The sensitivity of both *S*_*ητ*_ and *S*_*τη*_ was the lowest for RLT Rd II across all designs.

#### 2.4.1. RLT Rd I exhibits a wider amplitude range yet with a strong non-linear amplitude and period dependency

Numerical simulations of the RLT Rd I (Fig. 3B, E) indicate that the range of achievable amplitudes is increased by approximately 3.5-fold compared to RLT, yet with large period shifts. Specifically, the cross-sensitivity of period to amplitude S_*τη*_ is significantly larger compared to DFO Rd I (Fig. 2E). The shape of the tuneable region also indicates strong and highly non-linear coupling of the period with the amplitude. Increasing the amount of the protease *C* results in shorter periods but with a strong coupling between amplitude and period. Indeed, the range of achievable periods was narrower than in the original RLT but with an extended range for amplitude.

#### 2.4.2. RLT Rd II exhibits a wide range of periods and amplitudes tuneable in a near-independent manner

The simulations of RLT Rd II show that the amplitude is tuneable independently of the period as shown in Fig. 3C, F. Note that in this case, we exploit the architecture of RLT to tune the period by introducing a dual-input promoter at the second position of the ring. This position was chosen as it is not directly regulating or being regulated by the position of the output (G) and is expected to have the least possible period to amplitude sensitivity (S_*τη*_). The latter is further supported by simulation results shown in SI (S5). Simulation results indicate a significantly wider range of tuneable periods. More importantly, tuning the period results in low cross-sensitivity S_*ητ*_ (period to amplitude variations). This follows from the fact that the duration of the cycle of each repressor affects the overall period, but not the amplitude of oscillations of the other repressors. In fact, this effect becomes more apparent if more genes are added in the ring, as seen in simulation results of the generalised repressilator^32-34^ (with *n =* 5 and *n = 7* repressors) in SI (Fig. S5).

### 2.5 Application of the re-designed RLT Rd II: An orthogonally tuneable oscillator as a multiple-input single-output biosensor

Orthogonally tuneable oscillators can be used to precisely regulate the magnitude and temporal characteristics of downstream networks or metabolic pathways^1,3,11^. An alternative application is to reverse their use: instead of applying specific inputs to generate periodic oscillations of desired amplitude and period, the user can read the periodic signal and accurately determine the concentrations of the inputs; hence using the oscillator as a *dual-input single-output biosensor*^35^. According to our computational analyses, the RLT Rd II re-design is most orthogonal in terms of its cross-sensitivities, and would thus be most appropriate to build such a biosensor. The inputs can be mediated by any transcriptional regulatory network, including TFs, two component systems, or even CRISPR derived circuits^36,37^, that respond to ligands of interest (*e.g.* arsenic^38^ or aspartate^39^) or environmental signals (*e.g.* light^40-42^ or temperature^43^).

Fig. 4 illustrates the concept with RLT Rd II used as a biosensor. The network is exposed to ligand 1 (*I*_*1*_), which leads to modulation of the amplitude, and to ligand 2 (*I*_*2*_), which modulates the period of the oscillations. The output reporter (*e.g*., green fluorescent protein (GFP)) can be recorded with a basic setup comprising a light excitation source and a sensor that records the periodic fluorescent output over time. The analysis of the signal would map the period to the level of *I*_*2*_ using a precomputed function (red line); for a given *I*_*2*_, the amplitude versus *I*_*1*_ precomputed function then returns the concentration of *I*_*1*_.

**Figure 4.**
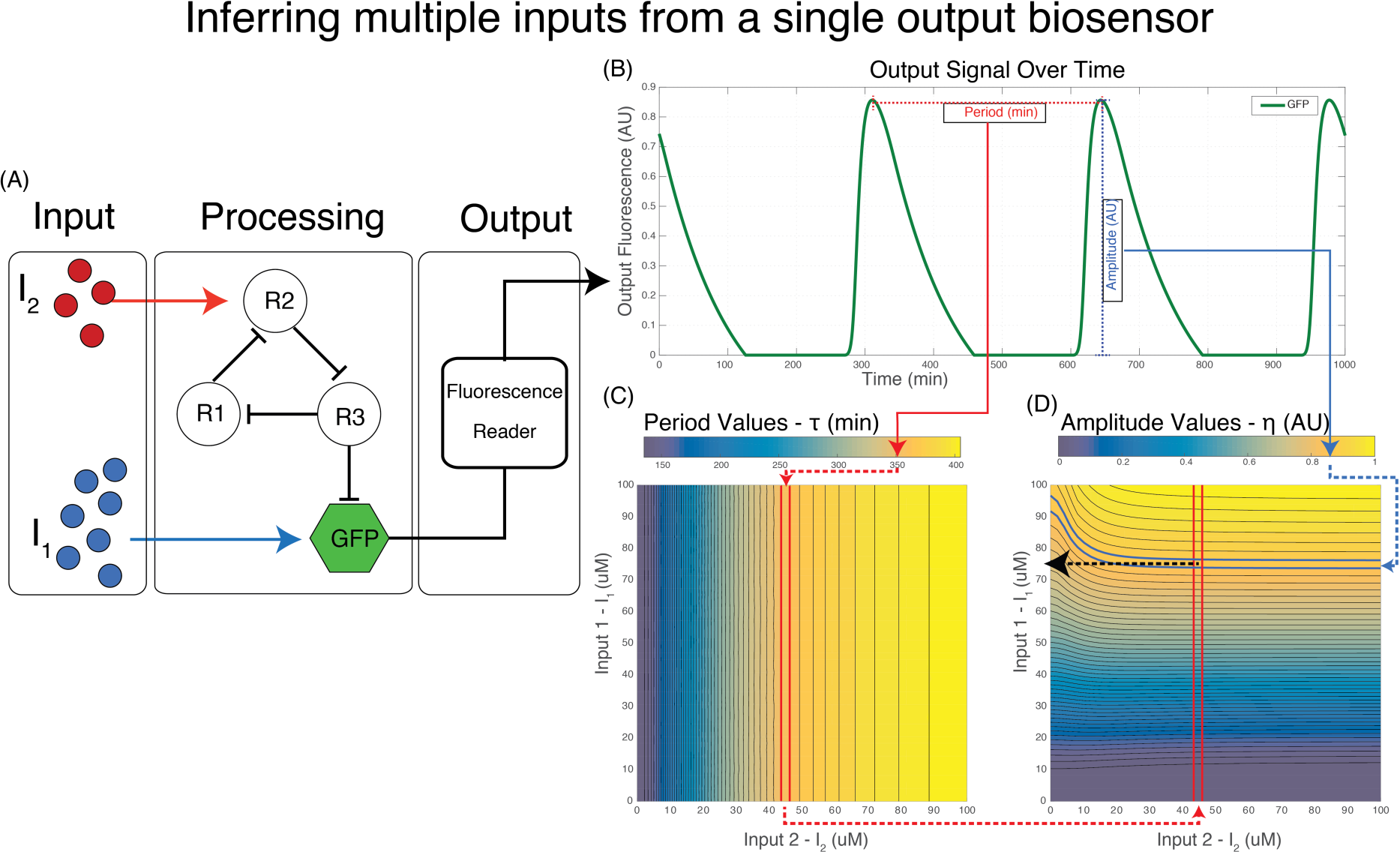
Diagram of a conceptual two input, single output oscillator based biosensor. **(A)** A diagram of a hypothetical biosensor where the concentration of two small molecules of interest *I*_*1*_ and *I*_*2*_ determine the characteristic of the oscillator in the processing unit which generates an oscillatory fluorescence signal, read in the output module. **(B)** The readout of the biosensor is analysed over a period to determine the amplitude and period of the oscillation. **(C)** Contour map showing the period as a function of both inputs. The isoclines are vertical as *I*_*1*_ has no effect on the period, therefore at this stage the concentration of *I*_*2*_ is determined by mapping the period to a standard curve of period versus *I*_*2*_. **(D)** Contour plot of the amplitude versus both inputs. Once the *I*_*2*_ concentration is determined from (B) the *I*_*1*_ concentration is simply found at the intersection point of the isoclines bounding the measured amplitude value (blue isoclines) and the isocline of *I*_*2.*_ The error for *I*_*1*_ and *I*_*2*_ is determined by the width of the blue and red isoclines, respectively.

This approach could have significant advantages over co-expressing multiple single – input – single-output biosensors. A multi – input – single-output biosensor requires a single excitation source and sensor, and a simpler electronic interface for implementing a low-cost biosensor. From a biological perspective, a single reporter imposes less metabolic burden on the host, and periodic expression of a single fluorescent reporter would reduce the toxic effects, *e.g.* due to production of free radicals upon excitation^44^. Finally, the error due to stochastic gene expression effects and measurement method, can be reduced in periodic signals by increasing the duration of the readings, in contrast to the single readings from traditional biosensors.

## 3. Discussion

Using computational modelling, this work has identified network re-designs that expand the range of achievable amplitudes and periods of the two oscillators most widely used in Synthetic Biology, the DFO and RLT. Additionally, we have identified those re-designs that enable orthogonal tuning of amplitude and period. In particular, we have shown that the RLT outperforms the DFO in achieving independent tuning of amplitude and period and extending the range of their achievable values. For instance, our DFO redesign showed fast oscillations (period below 1 hour) but within a limited range of achievable amplitudes compared to the RLT. This is due to the fact that the activating and repressing TFs in the DFO oscillate in phase, while in the RLT each repressor is expressed in succession such that the effects of tuning a repressor are temporarily separated to some extent.

*In vivo* implementations of the system would be affected by other couplings, such as competition for shared cellular resources (*e.g.,* energy, amino acids, ribosomes and polymerases^45,46^). This means that the range over which amplitude and period can be tuned independently should become narrower for experimental conditions where the TFs or output gene are overexpressed. To attenuate host context effects, the oscillator TFs should be expressed at low enough levels, *i.e.* levels at which burden and resource competition effects are not dominant. Recently, TX-TL implementation of these oscillators has been successfully demonstrated in cell-free microfluidic based platforms^47^ where a continuous flow of TX-TL^48^ rich in nutrients, translational and transcriptional resources was used. In this setup, the competition for shared resources is minimised and thus we anticipate this setup to constitute the best platform for a successful implementation of our re-designs.

A successful implementation of an orthogonally tuneable oscillator, such as our proposed RLT Rd II, would offer a powerful basis for several applications including our proposed dual-input single-output biosensor. It could also be used to generate periodic inputs to drive downstream networks, which can be useful to better understand the dynamics of cellular and biological systems^21,22,49,50^. Furthermore this can be utilised to optimise temporal enzyme expression in metabolic pathways^22^ or even pulsatile (*e.g.* circadian) release and administration of drugs and therapeutic molecules^51-53^.

### Engineering design principles of orthogonally tuneable oscillators with increased ranges of achievable amplitude and period

Through the computational simulation and analysis that we have carried out above and further data provided in the Supplemental Material, we identify the following core principles for the implementation of an orthogonally tuneable oscillator:

**a) Orthogonality of the degradation of the output with respect to the degradation of the oscillator’s TFs enables decoupling of the amplitude and period tuning.** In our proposed re-designs (DFO Rd II and RLT Rd II) this was realised by considering a second orthogonal protease pathway dedicated to the output. Alternatively, this decoupling can be achieved by removing the enzymatic degradation altogether like it was recently demonstrated experimentally for the repressilator^54^. The trade-off for using the latter approach is that the achievable periods will inevitably become longer and dependent on the growth rate of the host.

**b) Modulation of the output expression rate independently of the oscillator’s TF levels is required for setting the amplitude levels.** We simulated the above by considering a dual-input promoter that oscillates due to the periodic expression of the oscillator’s repressors. The rate of transcription obtained from this dual-input promoter is tuneable through the use of its second input, which in our case was an activator orthogonal to the oscillator’s repressors. Nevertheless, the same effect can be obtained by modulating other gene expression elements like the promoter, RBS, gene copy number or the degradation tags. The downside is that this modulation will require the creation of different genetic constructs for a required amplitude level and is not easily or externally tuneable in a dynamic manner.

**c) Control over the degradation rate of the oscillator’s TFs enables period modulation.** This was demonstrated in DFO Rd II where the protease abundance is controlled by an orthogonal input which, in turns, regulates the amount of activating TF (Y). Again, alternatively the same effect can be obtained by varying the protease levels through the use of promoters, RBSs and degradation tags of different strengths or by varying the copy number of the protease gene. However, when doing so, the range of achievable amplitudes was observed to be much more dependent on protease abundance (Fig. 2F, Fig. S3D).

**d) Varying the strength of the negative feedback loop enables control of the period of the oscillations.** As we showed in our re-designs, controlling the expression rate of one of the repressilator’s repressors in RLT Rd II by the use a dual-input promoter enables period modulation. This promoter retains the original regulation from the oscillator’s TF but in addition is also responsive to an orthogonal activating TF (Y). As mentioned above, modifying the gene expression elements (promoter, RBS, etc.) of the affected repressor will generate the same effects. However, when doing so for the DFO case, the range of achievable periods was significantly narrower (Fig 3F, Fig S3B.)

**e) Increasing the network distance between the modulated repressor and the RLT allows for nearly completely orthogonal period to amplitude modulation.** When we considered repressilator rings with 5 and 7 repressors our models predicted that, when the rate of expression of one of the repressors was tuneable as per point d), the period had the least effect on the amplitude when the repressor was not directly regulating or regulated by the output (SI, S5).

## 4. Computational Methods

The models given in section 3 were computationally implemented in MATLAB’s Simbiology V3 as a set of reactions, species, parameters and rules (Tables given in SI Tables 7.1-7.8). The models were simulated using the ODE15s (Stiff / NDF) solver. The simulated time for individual runs was set to 1000 min in the case of the DFO and its redesigned models and 4000 min in the case of the RLT and its corresponding redesigns. The RLT required longer simulations time to be able to capture a sufficient number of oscillations, which is important for downstream analysis as the RLT produced longer periods (150-400 min) compared to the DFO (15-55 min) as shown in Fig 3. Each input was parameter scanned from 0 to saturating amounts of input (100 uM) in 50 steps. Each simulation dataset was obtained through parallel computations of the 50 X 50 dual-input titration matrix associated with these parameter scans.

The simulation results were analysed using custom-written scripts in MATLAB. The analysis was performed by, first, extracting the trajectories of the output (*G)* and the transcription factors. This was then followed by identification of the peaks of the signal using the ‘findpeaks’ function of MATLAB. The period was then calculated by determining the time difference for consecutive peaks. The amplitude was determined by computing the difference between the peak and minimum levels of *G* for a given cycle.

